# Ultra-fast genetic recovery of dead fish through cross-family germline stem cell transplantation

**DOI:** 10.64898/2026.01.05.697817

**Authors:** Chaofan Wang, Huiying Ma, Xiaosi Wang, Yongkang Hao, Junwen Zhu, Houpeng Wang, Yaqing Wang, Xiaxia Gao, Mudan He, Songlin Chen, Yonghua Sun

## Abstract

The conservation of genetic resources from aquaculture species and endangered fish is increasingly challenged by large body size, long reproductive cycles, and limited opportunities for timely intervention after death. Here, we establish and validate an ultra-fast genetic platform based on germline stem cell transplantation to enable postmortem genetic recovery in fish. Using grass carp (*Ctenopharyngodon idella*) as a representative warm-water species, we systematically quantified the relationships among postmortem tissue freshness, germline stem cell viability, and transplantation efficiency, and demonstrated that low-temperature preservation plays a decisive role in maintaining germline activity after death. Germline stem cells isolated from deceased grass carp were transplanted into germ-cell–depleted zebrafish recipients, where they rapidly colonized recipient gonads, underwent proliferation and differentiation, and generated functional donor-derived gametes within three months. These gametes supported successful fertilization and normal embryonic development, ultimately yielding viable grass carp offspring. Our results reveal an intrinsic postmortem resilience of germline stem cells and demonstrate that cross-species transplantation into small, fast-maturing hosts can dramatically accelerate genetic recovery. This strategy overcomes key biological and logistical constraints associated with conventional breeding-based rescue approaches and provides a rapid, scalable, and broadly applicable framework for postmortem genetic resource conservation in aquaculture and endangered fish species.

## 1. Introduction

Fish breeds carrying superior genetic traits are the core resources of aquaculture. However, under captive conditions, the risks of pathogen infection and facility failure are ever-present. Once an accident occurs, it may not only lead to the loss of valuable cultured individuals but also cause the permanent disappearance of precious genetic resources in economically important fish species. Similarly, the unexpected death of endangered fish can result in the irretrievable loss of their rare genetic heritage (Li, 2022; Lind *et al*, 2012). Therefore, it is urgently necessary to establish a reliable and feasible technological strategy to achieve the genetic revival of deceased fish, thereby maximizing the preservation and inheritance of essential aquatic genetic resources.

Germline stem cells (GSCs) serve as the fundamental carriers for transmitting genetic information across generations. In recent years, GSC transplantation–based surrogate reproduction has emerged as an important technology in fish genetic breeding (Gui *et al*, 2022; Houston *et al*, 2020). Functional donor-derived sperm production has been reported in multiple fish species through transplantation into surrogate hosts (Yoshizaki & Yazawa, 2019). For example, this technique has enabled surrogate reproduction between Pacific salmon and rainbow trout, allowing repeated production of salmonid gametes (Yoshizaki *et al*, 2024). In addition, spermatogonial stem cell transplantation has achieved cross-subfamily surrogate reproduction between juji (*Gobiocypris rarus*) (Su *et al*, 2025) and zebrafish (*Danio rerio*), successfully generating gene-edited sperm derived from a donor species across subfamily boundaries (Wang *et al*, 2023; Zhang *et al*, 2022a). More notably, a recent study demonstrated ultrafast production of functional grass carp (*Ctenopharyngodon idellus*) gametes in zebrafish hosts via GSC transplantation (Sun *et al*, 2024). This breakthrough suggests that GSC transplantation may overcome limitations imposed by differences in body size, husbandry mode, reproductive characteristics, and even phylogenetic distance. Thus, by leveraging small fish with short maturation cycles and facile breeding, it may become feasible to rapidly resurrect the genetic lineage of large-bodied, slowly maturing, and breeding-challenging fish that have died. Furthermore, modifying the immune system of transplantation recipients can markedly enhance the efficiency of surrogate reproduction (Kawasaki *et al*, 2016), raising the possibility of achieving ultrafast genetic revival across even wider evolutionary distances.

Despite these advances, standardized procedures for isolating viable GSCs from deceased fish for genetic revival are still lacking. Previous studies have shown that type A spermatogonia (ASGs) isolated from dead rainbow trout can survive, proliferate, and differentiate after intraperitoneal transplantation into live conspecific recipients, thereby establishing a preliminary framework for genetic restoration (Yang *et al*, 2022). However, functional gametes derived from dead donors have not yet been obtained, and thus true genetic resurrection of deceased individuals has not been achieved. Additionally, this approach has only been validated in intraspecific transplantation, and whether ultrafast genetic revival of deceased fish can be achieved across species boundaries remains unclear. More importantly, rainbow trout is a typical cold-water species, and criteria derived from gonadal deterioration in this species may not be directly applicable to warm-water fish, such as grass carp. Therefore, dedicated studies are urgently required to enable the genetic recovery of deceased individuals in warm-water aquaculture species.

In this study, we used grass carp as a model to systematically evaluate the freshness and GSC viability at different postmortem time points. We further assessed the colonization, proliferation, and differentiation potential of GSCs isolated from deceased grass carp by transplanting them into zebrafish recipients. By establishing the relationship between postmortem freshness and transplantation success, we ultimately achieved the genetic recovery of deceased grass carp individuals, remarkably completing this process within only three months by using zebrafish as a rapid surrogate host.

## 2. Materials and methods

### 2.1. Experimental Animals

Grass carp and zebrafish used in this study were obtained from the National Aquatic Biological Resource Center and the China Zebrafish Resource Center (NABRC and CZRC, Wuhan, China). Grass carp, approximately six months old, were maintained in freshwater at 26 °C. Wild-type AB strain zebrafish were maintained at 28 °C under a 14 h light/10 h dark photoperiod. Fertilized embryos were collected by natural spawning and subsequently used for microinjection.

### 2.2. Preparation of Dead Grass Carp Samples

Forty grass carp were randomly assigned to eight groups (n = 5 per group) and euthanized by immersion in an overdose of tricaine methanesulfonate (MS-222; 500 mg/L) for 15 min, followed by 10 min in running water to confirm death. Fish were then maintained either in running water at 26 °C for 0, 6, or 12 h post-death (hpd), or in an ice–water mixture (2–4 °C) for 0, 6, 12, 24, or 48 hpd. Prior to gonadal fixation and donor cell preparation, total length and body weight were recorded. The grass carp used in this study showed no significant differences in body size among groups (mean length 12.59 ± 1.59 cm; mean weight 31.51 ± 2.78 g; n = 5).

### 2.3. Evaluation of Freshness in Dead Grass Carp

Freshness of grass carp in each group was assessed by measuring gill color saturation, muscle ATP concentration, and whole-body rigor index. Gill images were captured under standardized lighting and color saturation was quantified using ImageJ. Muscle ATP levels were determined from ∼20 mg tissue samples using a commercial ATP assay kit (S0026, Beyotime, China), with luminescence measured by a luminometer (GloMax™ 20/20, Promega, USA) and calculated from a standard curve.

Rigor index was evaluated using the horizontal displacement method (Iwamoto *et al*, 1987). Fish were positioned horizontally with the posterior body extending beyond the table edge, and the vertical displacement of the tail was measured. Rigor index was calculated as R = [(D₀ − D) / D₀] × 100, where D₀ and D represent tail displacement before rigor and at measurement, respectively (Bito, 1983). Measurements were performed at 0, 6, and 12 h post-death for room-temperature groups and at 0, 6, 12, 24, and 48 h post-death for ice–water groups.

### 2.4. Integrity and Viability of Gonadal Cells in Dead Grass Carp

For cell dissociation, ∼20 mg gonadal tissue was digested in a mixed enzyme solution containing 0.25% trypsin (C0201, Beyotime, China), 300 U/mL collagenase (C128711, Aladdin, China), and 0.05% DNase I (04716728001, Roche, Switzerland) at 33 °C for 2 h with intermittent gentle pipetting. Cell suspensions were filtered (40 µm), washed with D-PBS containing 1% FBS (A5256701, Gibco, USA), and stained with acridine orange/propidium iodide (AO/PI). Gonadal cell viability was quantified using an automated cell counter (C200FL, Ruiwo, China), with three fields analyzed per sample.

For immunofluorescence, PFA-fixed tissues were permeabilized, blocked, and incubated overnight at 4 °C with mouse anti-Caspase-3 (31A1067, SCBT, USA) and rabbit anti-Ddx4 primary antibodies (purified by Tianda Scientific, Wuhan, China) (1:500). Samples were then incubated with Alexa Fluor–conjugated secondary antibodies, counterstained with DAPI, and imaged using a fluorescence microscope (Ye *et al*, 2023).

### 2.5. Preparation of Grass Carp Germ Cell Suspensions

The enzymatically dissociated cell suspension was processed following the reference method (Zhang *et al*., 2022a), using Percoll gradient centrifugation to enrich germ cells. Cells located between the 30%-40% Percoll gradient layers were collected, washed with D-PBS containing 1% fetal bovine serum (FBS), and resuspended. An aliquot of 5 µL was used for Nanos2 immunofluorescence staining, while the remaining cells were labeled with CellTracker CM-DiI and finally resuspended in 10 µL of L-15 medium supplemented with 1% FBS.

### 2.6. Preparation of Zebrafish Recipients

Zebrafish embryos were injected with 100 μM of *dnd end 1* (*dnd1*) morpholino oligonucleotide (*dnd1*_MO: 5′-GCTGGGCATCCATGTCTCCGACCAT-3′) to eliminate endogenous primordial germ cells. Injected embryos were cultured for 4-5 days in 0.3× Danieau’s buffer, and were used as recipients for germ cell transplantation.

### 2.7. Germ Cell Transplantation

Cell transplantation was performed under a stereomicroscope using a glass capillary needle (∼50 µm tip). Zebrafish larvae were anesthetized with MS-222 and positioned on 2% agarose plates. Approximately 30–50 donor cells were transplanted into the peritoneal cavity adjacent to the genital ridge, following a previously described method (Zhang *et al*., 2022a). After transplantation, larvae were recovered in 0.3× Danieau’s buffer at 28.5 °C and returned to the rearing system the next day. Germ cell colonization was assessed at 5 days post-transplantation by fluorescence microscopy.

### 2.8. Correlation analysis of postmortem freshness, gonadal cell viability, and post-transplantation germ cell colonization rate

Pearson correlation analysis was performed to evaluate the associations between gill color saturation, ATP concentration, rigor index, gonadal cell viability, and germ cell colonization rate. Univariate correlations were visualized using heatmaps. To determine whether colonization rate could be estimated using a readily observable parameter, univariate linear regression models were constructed for gill color saturation and gonadal cell viability, respectively, and the corresponding regression equations were derived. All statistical analyses and graphical visualizations were performed in R (version 4.3.3).

### 2.9. Germ Cell Proliferation and Subsequent RT-PCR Detection

To assess proliferation of transplanted grass carp germ cells in zebrafish recipients, EdU labeling was performed. Fifteen zebrafish at 30 days post-fertilization were incubated in 400 μM EdU for 48 h, followed by gonadal dissection and fixation in 4% PFA. EdU (C0078S, Beyotime, China) incorporation was detected using a commercial kit according to the manufacturer’s instructions.

Immunofluorescence staining was conducted on gonadal samples using anti-Ddx4 antibodies or pre-labeled fluorescent antibodies, with all buffers supplemented with ribonucleoside–vanadyl complex. Ddx4-positive gonads were subsequently subjected to RNA extraction, reverse transcription, and PCR analysis using standard protocols and, all primers used are listed in Supplementary Table S1.

### 2.10. Collection and Analysis of Recipient Zebrafish Sperm

Three months after transplantation, sperm samples were collected from zebrafish recipients and subjected to PCR identification or field emission scanning electron microscopy (FESEM) analysis. For FESEM, sperm were fixed in 2% glutaraldehyde, applied to poly-L-lysine–coated coverslips, dehydrated through a graded ethanol series, air-dried, platinum-coated, and examined using a FESEM system (S-4800, Hitachi, Japan). Sperm head and tail lengths were quantified using ImageJ, and statistical analyses were performed using unpaired two-tailed Student’s t-tests, with data presented as mean ± SD.

### 2.11. In vitro fertilization using recipient-derived sperm and whole-genome resequencing of offspring

Gonads from germline chimeric zebrafish recipients were dissected into 500 µL Hank’s buffer and minced into small fragments using fine scissors to release sperm. Approximately 200 µL of sperm suspension was mixed with ∼1 mL of grass carp eggs and gently agitated. After incubation for several minutes, excess sperm were removed by rinsing with clean water. Fertilization rate was assessed at 6 h post-fertilization by counting developing embryos, and hatched larvae were subsequently monitored. To evaluate the genetic integrity of offspring derived from germ cell transplantation, larval samples were collected for whole-genome resequencing analysis.

## 3. Results

### 3.1 Evaluation of Postmortem Freshness in Grass Carp

Because the freshness of dead fish rapidly deteriorates at room temperature, we first conducted a systematic evaluation of postmortem freshness in grass carp under low-temperature conditions using an ice-water mixture. In aquaculture practice, gill color is commonly used as an intuitive indicator to estimate postmortem time (Shi *et al*, 2018). Immediately after death, grass carp gills exhibit a bright red coloration; however, with increasing postmortem time, the gill color gradually fades and ultimately becomes grayish-white with evident tissue degradation (Fig. 1A). Quantitative analysis of gill red color saturation using ImageJ revealed a significant linear decline over time (Fig. 1B). We next measured ATP levels in muscle tissues at different postmortem time points. Muscle ATP concentrations decreased rapidly within the first 6 h post-death (hpd) and became almost undetectable by 24 hpd (Fig. 1C). In parallel, assessment of the rigor index showed that body stiffness progressively increased with postmortem time, and fish had not yet entered the resolution phase of rigor mortis even at 48 hpd (Fig. 1D). Collectively, these results indicate that, even under low-temperature conditions, the overall freshness of grass carp declines markedly within 12 hpd.

**Figure 1.**
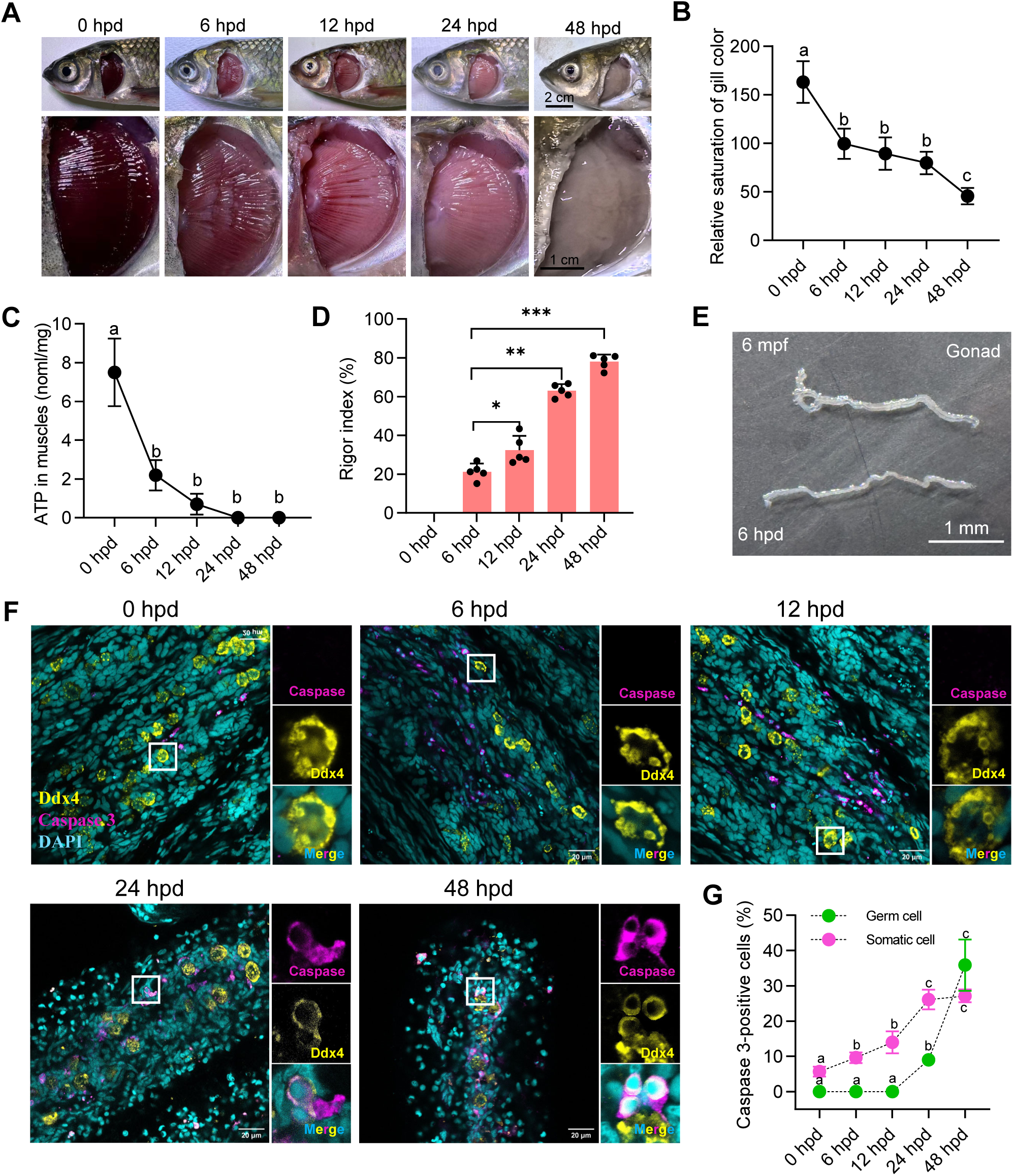
Physical and biochemical characteristics of grass carp at different times post-death during ice-water storage and apoptosis of gonadal germ cells. A: Representative images of gills from grass carp at 0, 6, 12, 24, and 48 hpd. B: Gill color saturation at different hpd. C: Muscle ATP content at different hpd. D: Rigor index at different hpd. Data are presented as mean ± SEM (N = 5). Different letters and asterisks indicate significant differences (p < 0.05, Tukey–Kramer test). E: Representative gonadal morphology of 6-month-old grass carp. F: Representative images of *Caspase-3* and *Ddx4* double immunofluorescence staining in gonads at different hpd. *Ddx4* marks germ cells, *Caspase-3* marks apoptotic cells, and nuclei are counterstained with DAPI. G: Proportion of *Caspase-3*-positive germ cells at different hpd. Data are presented as mean ± SEM (N = 3). Different letters indicate p < 0.05 (Tukey–Kramer test).

To further assess the viability of germ cells in the gonads, double immunofluorescence staining was performed using antibodies against Caspase-3 and Ddx4. Caspase-3 signals were detected in gonadal tissues preserved in ice water at 0, 6, 12, 24, and 48 hpd. Notably, no Caspase-3/Ddx4 double-positive cells were observed at 0, 6, or 12 hpd. At 24 hpd, a subset of Ddx4-positive germ cells began to exhibit Caspase-3 positivity, and the number of double-positive cells further increased at 48 hpd (Fig. 1F). In contrast, Caspase-3 signals in somatic cells were detectable as early as 0 hpd and progressively increased over time. Quantitative analysis revealed that the proportion of Caspase-3–positive germ cells was significantly elevated at 48 hpd and exceeded that observed in somatic cells (Fig. 1G). These findings indicate that, under low-temperature conditions, germ cells in the gonads retain viability for a longer period than somatic cells.

### 3.2 Colonization of Germline Stem Cells Isolated from Deceased Grass Carp in Zebrafish

To further evaluate the viability of germ cells from deceased fish, we performed live-cell assays and germline stem cell (GSC) transplantation experiments. GSC transplantation represents a key strategy for genetic recovery of dead individuals; however, preparation of recipient fish requires time, and GSC transplantation is constrained by a defined optimal time window in recipients. Therefore, immediate transplantation following death is not feasible. To address this limitation, we employed an established long-term cryopreservation and recovery protocol for fish gonadal cells (Wylie *et al*, 2025). Gonads isolated from deceased fish were cryopreserved for 24 h, subsequently thawed, and used for GSC transplantation (Fig. 2A). Single-cell suspensions obtained from thawed gonadal tissues were first stained with propidium iodide (PI) to assess cell viability. The proportion of PI-positive cells increased progressively with postmortem time, with a pronounced increase observed in samples collected at 48 hpd) (Fig. 2B, C). Cells from the 30–40% Percoll density gradient fractions were collected and subjected to immunofluorescence staining using a laboratory-generated anti-*nanos2* antibody to evaluate the enrichment efficiency of germline stem cells following Percoll centrifugation (Fig. 2D) (Wang *et al*, 2022). The proportion of *nanos2*-positive cells was only 5.37 ± 1.7% in samples without Percoll centrifugation, whereas Percoll-enriched cell suspensions exhibited a markedly higher *nanos2*-positive rate of 33.2 ± 6.3%, indicating that Percoll density gradient centrifugation effectively enriched germline stem cells (Fig. 2E).Germline stem cells were then enriched from gonadal single-cell suspensions derived from different postmortem time points using a Percoll density gradient centrifugation method established in our laboratory (Sun *et al*., 2024; Zhang *et al*., 2022a). The isolated GSCs were transplanted into zebrafish hosts in which endogenous germ cells had been ablated, and colonization efficiency was quantified.

**Figure 2.**
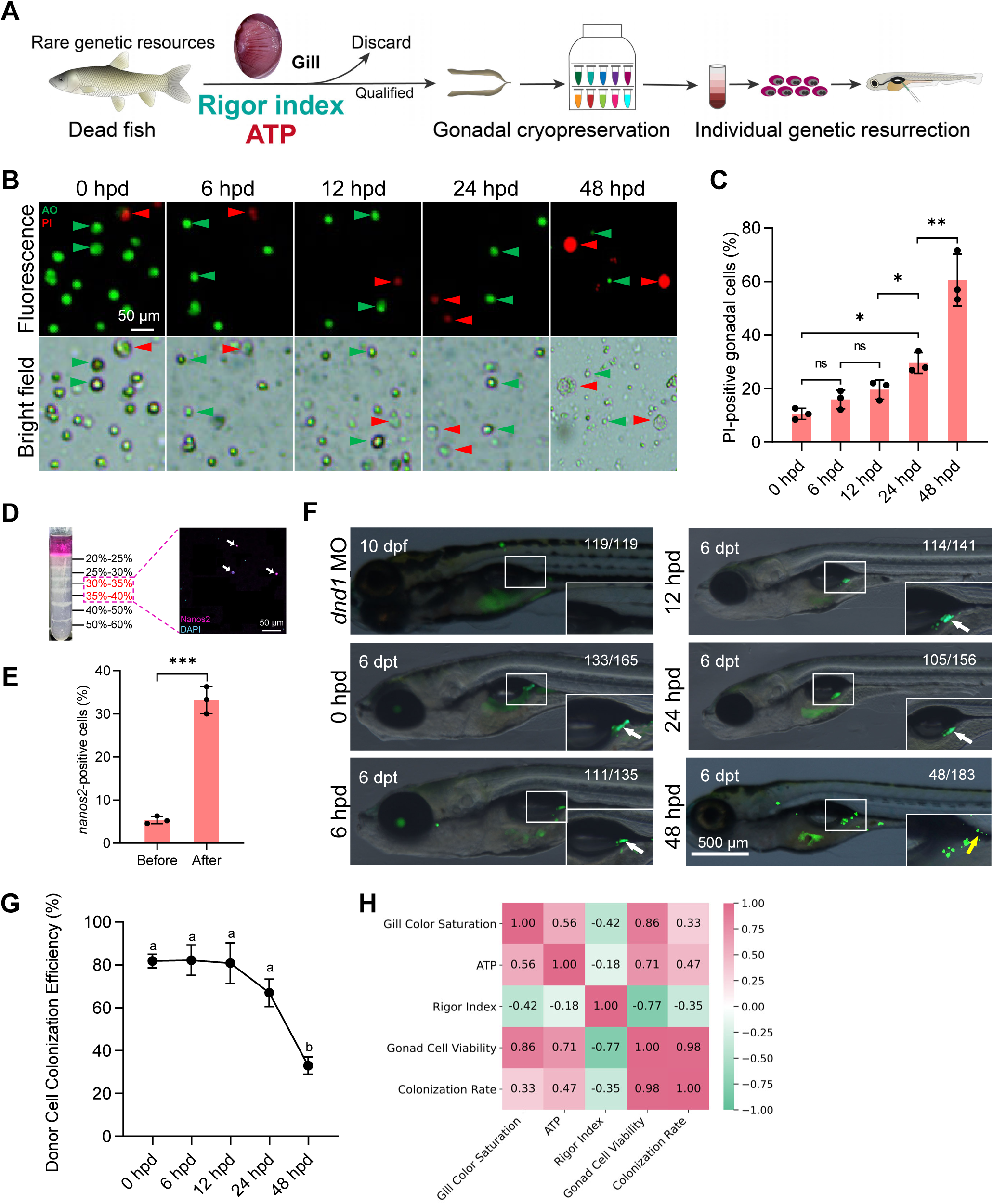
Viability, enrichment, transplantation efficiency of postmortem-derived gonadal cells, and correlation with colonization rate. A: Schematic overview of postmortem handling, preservation, and genetic reconstruction of valuable fish germplasm. B: AO/PI staining of recovered gonadal single-cell suspensions at 0, 6, 12, 24, and 48 hpd. C: Percentage of PI-positive cells at different hpd (mean ± SEM, N = 3). ns, not significant; p < 0.05 (Student’s t-test). D: Immunofluorescence staining of cells collected from the 30–40% Percoll density fractions showing *nanos2*-positive germline stem cells. E: Proportion of *nanos2*-positive cells among total cells (p < 0.05, Student’s t-test). F: Fluorescent images showing donor-derived germ cells colonized in the gonadal ridge of zebrafish hosts (arrows). The yellow arrow represents an incomplete cell. G: Colonization rates of donor germ cells derived from grass carp at different hpd (mean ± SEM, N = 3). Different letters indicate p < 0.05 (Tukey–Kramer test). H: Heatmap showing Pearson correlations between freshness-related variables, gonadal cell viability, and colonization rate.

At 6 h post-transplantation, GSCs isolated from fish at 0, 6, and 12 hpd exhibited colonization rates exceeding 70% in zebrafish recipients. GSCs derived from fish at 24 hpd showed a moderate reduction in colonization efficiency (68.5 ± 8.5%), which was not statistically different from the 0 hpd group. In contrast, GSCs isolated at 48 hpd displayed a highly significant reduction in colonization efficiency, reaching only 34.5 ± 4.1% (Fig. 2D). Notably, these cells exhibited blurred cellular boundaries and abundant cellular debris, indicating compromised cellular integrity (Fig. 2D, E). Together, these results demonstrate that GSC viability declines markedly at 48 hpd, impairing their ability to efficiently colonize and survive in host gonads. This finding is consistent with earlier postmortem freshness assessments based on gill color saturation and gonadal apoptosis levels.

Furthermore, to investigate the relationship between fish freshness indicators—including gill color saturation, ATP content, rigor mortis index, and gonadal cell viability—and germline stem cell colonization efficiency, we performed correlation analyses (Fig. 2F). The results showed that gill color saturation was moderately positively correlated with colonization efficiency (R = 0.33), whereas ATP content exhibited a stronger positive correlation (R = 0.47). In contrast, the rigor mortis index was negatively correlated with colonization efficiency (R = −0.35). Notably, gonadal cell viability displayed the strongest correlation with successful germline stem cell colonization, with a correlation coefficient as high as 0.98, indicating that cell viability is likely the most critical predictor of transplantation potential in postmortem fish.

### 3.3 Effects of Postmortem Freshness Changes at Room Temperature in Grass Carp

Grass carp thrive within an optimal temperature range of 20-32 °C, under which postmortem deterioration proceeds rapidly. When maintained at room temperature (25 °C), a pronounced decline in gill color saturation was observed as early as 6 hpd, accompanied by the onset of gill filament whitening. By 12 hpd, the gills had completely lost their coloration, and the eyes became visibly opaque (Fig. 3A). Consistently, muscle ATP content declined sharply to below 0.5 nmol/mg within 6 hpd. Corresponding to this rapid ATP depletion, the rigor mortis index reached its maximum at approximately 6 hpd (Fig. 3B-C). By 12 hpd, muscle ATP levels dropped to 0 nmol/mg, and the rigor index began to decrease, indicative of the onset of rigor resolution (Fig. 3D). To further evaluate gonadal cell viability, postmortem fish were dissected and gonadal single-cell suspensions were analyzed. At room temperature, a dramatic increase in the proportion of PI-positive cells was detected at 6 hpd, indicating a substantial loss of gonadal cell viability (Fig. 3E, F). Notably, by 12 hpd, severe decomposition of the abdominal cavity precluded the isolation of intact gonadal tissue (Fig. 3E). Collectively, these observations demonstrate that, for warm-water fish species, genetic recovery of deceased individuals necessitates immediate cold storage following death.

**Figure 3.**
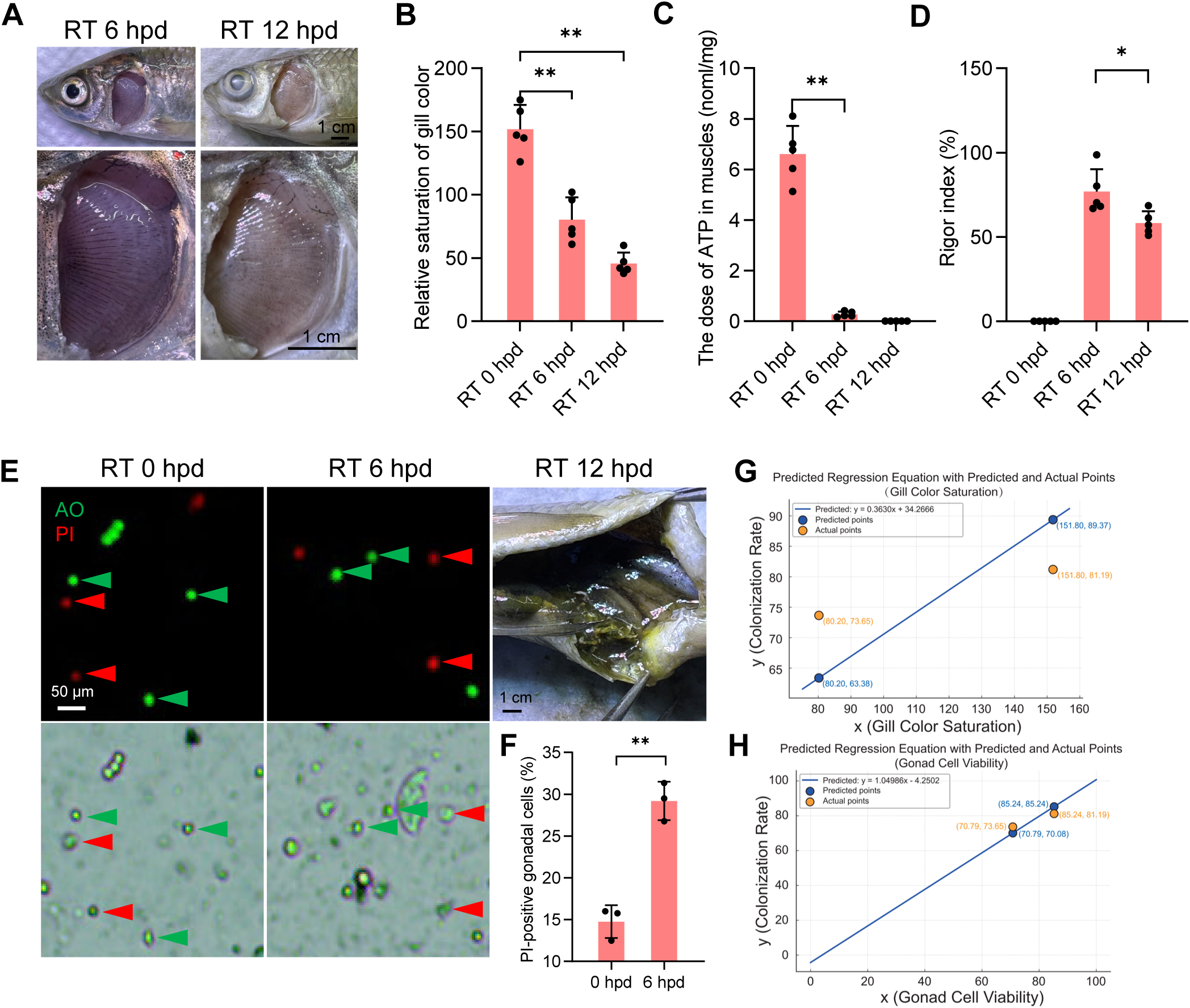
Freshness parameters of grass carp stored at room temperature and prediction of colonization rate. A: Representative gill images at 6 and 12 hpd. B: Gill color saturation at different hpd. C: Muscle ATP content at different hpd. D: Rigor index at different hpd. E: AO/PI staining of recovered gonadal single-cell suspensions at 0 and 6 hpd, and representative abdominal cavity image at 12 hpd. F: Percentage of PI-positive gonadal cells at different hpd. G: Univariate regression analysis and prediction of colonization rate based on gill color saturation. H: Univariate regression analysis and prediction of colonization rate based on gonadal cell viability. Data are presented as mean ± SEM (N = 5). p < 0.05 (Student’s t-test).

Given the rapid postmortem deterioration of grass carp at room temperature—characterized by precipitous ATP depletion and an early peak followed by resolution of rigor mortis—the regression models established under low-temperature conditions to link postmortem freshness indicators with germline stem cell colonization efficiency are not applicable at room temperature. Accordingly, we independently constructed regression models using gill color saturation and gonadal cell viability to predict colonization efficiency, and compared the predicted values with experimentally observed outcomes. At 0 h postmortem, the mean gill color saturation was 151.8, yielding a predicted colonization rate of 89.37%, whereas the actual colonization rate was 81.19%. At 6 hpd, gill color saturation decreased to 80.2, corresponding to a predicted colonization rate of 63.38%, compared with an observed rate of 73.65% (Fig. 3G). In contrast, predictions based on gonadal cell viability more closely approximated the actual colonization rates. Specifically, at 0 hpd, gonadal cell viability was 85.24%, resulting in a predicted colonization rate of 85.24%, closely matching the observed rate of 81.19%. At 6 hpd, gonadal cell viability was 70.79%, with a predicted colonization rate of 70.08%, again closely aligning with the observed rate of 73.65% (Fig. 3H). Although gonadal cell viability provides a more accurate predictor of post-transplantation germline stem cell colonization efficiency, gill color saturation remains a simple, rapid, and operationally practical indicator for on-site assessment under aquaculture conditions.

### 3.4 Proliferation and Differentiation of Germline Stem Cells Derived from Deceased Grass Carp in Zebrafish

To further characterize the growth and proliferative behavior of transplanted germ cells within host gonads, we labeled germ cells using an antibody against Ddx4 and assessed cell proliferation by EdU incorporation. Compared with the dnd1 MO–injected control group, zebrafish hosts transplanted with germ cells from all donor groups except the 48 hpd group exhibited restoration of germ cell populations in their gonads, with EdU-positive signals detected in multiple germ cells (Fig. 4A). Quantification of germ cell numbers and EdU-positive cells across three independent microscopic fields revealed no significant difference between hosts transplanted with germ cells from 0 hpd and 6 hpd donors. In contrast, the number of germ cells in hosts transplanted with 12 hpd donor cells was significantly reduced compared with the 6 hpd group, whereas no significant difference was observed between the 12 hpd and 24 hpd groups. Notably, no germ cells were detected in host gonads transplanted with germ cells derived from 48 hpd donors (Fig. 4B). In addition, robust EdU incorporation was observed in germ cells within recipient gonads transplanted with germ cells from 0, 6, 12, and 24 hpd donors. These results indicate that although the proportion of chimeric gonads containing germ cells gradually decreased with increasing postmortem time, germline stem cells derived from donors at 0, 6, 12, and 24 hpd retained biological activity and proliferative capacity in hosts.

**Figure 4.**
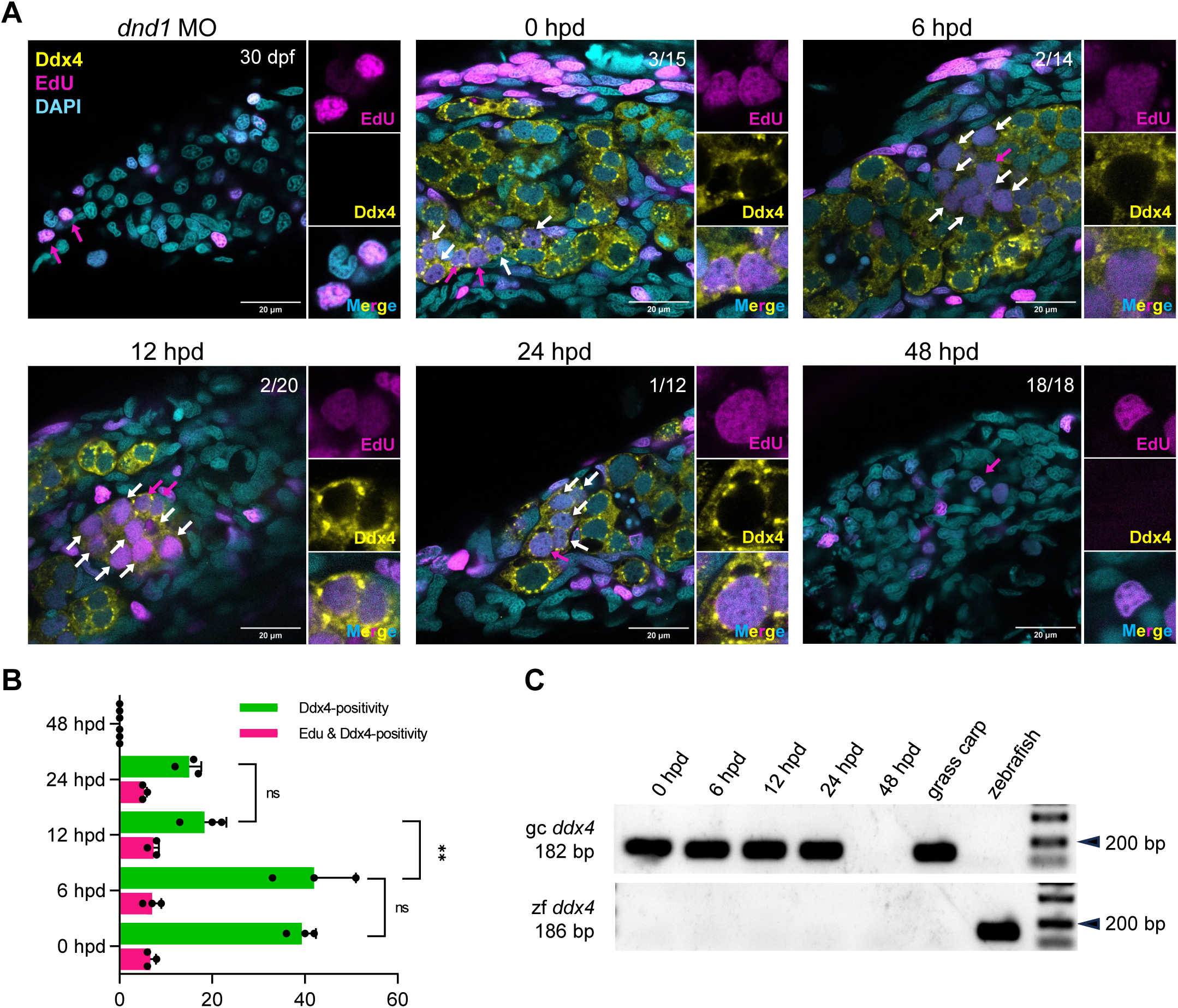
Proliferation of donor-derived germ cells in recipient gonads and correlation with freshness-related variables. A: Representative images of *Ddx4* and EdU double staining in recipient zebrafish gonads after transplantation. B: Quantification of donor-derived germ cells and EdU-positive cells in recipient gonads following transplantation of donor cells derived at different hpd (mean ± SEM, N = 3). ns, not significant; p < 0.05 (Student’s t-test). C: RT-PCR detection of gc___*ddx4* and zf___*ddx4* transcripts in host gonads.

To further verify the donor origin of germ cells in chimeric gonads, total RNA was extracted from recipient gonads and reverse-transcribed into cDNA. RT-PCR analyses were then performed using species-specific ddx4 primers for grass carp (*gc_ddx4*) and zebrafish (*zf_ddx4*). Specific amplification products were detected exclusively with the grass carp–specific gc_ddx4 primers in chimeric gonads transplanted with germ cells from 0, 6, 12, and 24 hpd donors, whereas no amplification was observed with zebrafish-specific zf_ddx4 primers (Fig. 4C). These results confirm that germ cells present in the chimeric gonads were derived solely from donor grass carp rather than from zebrafish. In contrast, no grass carp ddx4 transcripts were detected in host gonads transplanted with germ cells from 48 hpd donors, indicating the absence of donor-derived germ cells in this group. Collectively, these findings demonstrate that under low-temperature preservation conditions, germline stem cells isolated from grass carp within 24 h after death can survive, proliferate, and maintain germline identity in zebrafish hosts, whereas germline stem cells isolated at 48 h postmortem lose their capacity for survival and proliferation in host gonads.

### 3.5 Zebrafish Recipients Produce Functional Gametes Derived from Deceased Grass Carp

Three months after transplantation, zebrafish hosts were screened by gentle abdominal pressure to collect semen-like samples, which were subsequently subjected to PCR analysis. Two PCR-positive samples were detected in the 0 hpd group, whereas one PCR-positive sample was identified in each of the 6, 12, and 24 hpd groups (Fig. 5A). The statistical results revealed an overall declining trend starting from 0 hpd, with a significant reduction observed at 12 hpd. By 24 hpd, some experimental groups no longer yielded any PCR-positive semen samples, and no PCR-positive signals were detected in any group derived from fish deceased for 48 hpd (Fig. 5B). PCR-positive zebrafish recipients were subsequently dissected, and their gonads were examined under a stereomicroscope. In wild-type zebrafish, the testes appeared largely opaque, well developed, and densely packed with mature sperm. In contrast, gonads from hosts transplanted with grass carp germline stem cells exhibited extensive translucent regions, appeared relatively atrophic, and contained markedly fewer spermatozoa. As expected, testes from dnd1 MO–injected control zebrafish were completely transparent and devoid of sperm (Fig. 5C). Histological sectioning and immunofluorescence analyses were further performed. Ddx4 staining revealed abundant sperm cells, along with small numbers of spermatogonia and spermatocytes, in wild-type zebrafish testes. Similarly, chimeric gonads derived from recipients transplanted with grass carp germline stem cells also contained large numbers of sperm cells. In contrast, no germ cells were detected in the gonads of dnd1 MO–injected zebrafish (Fig. 5D). These results demonstrate that germline stem cells isolated from deceased grass carp gonads can differentiate and produce donor-derived gametes in zebrafish hosts within only three months post-transplantation.

**Figure 5.**
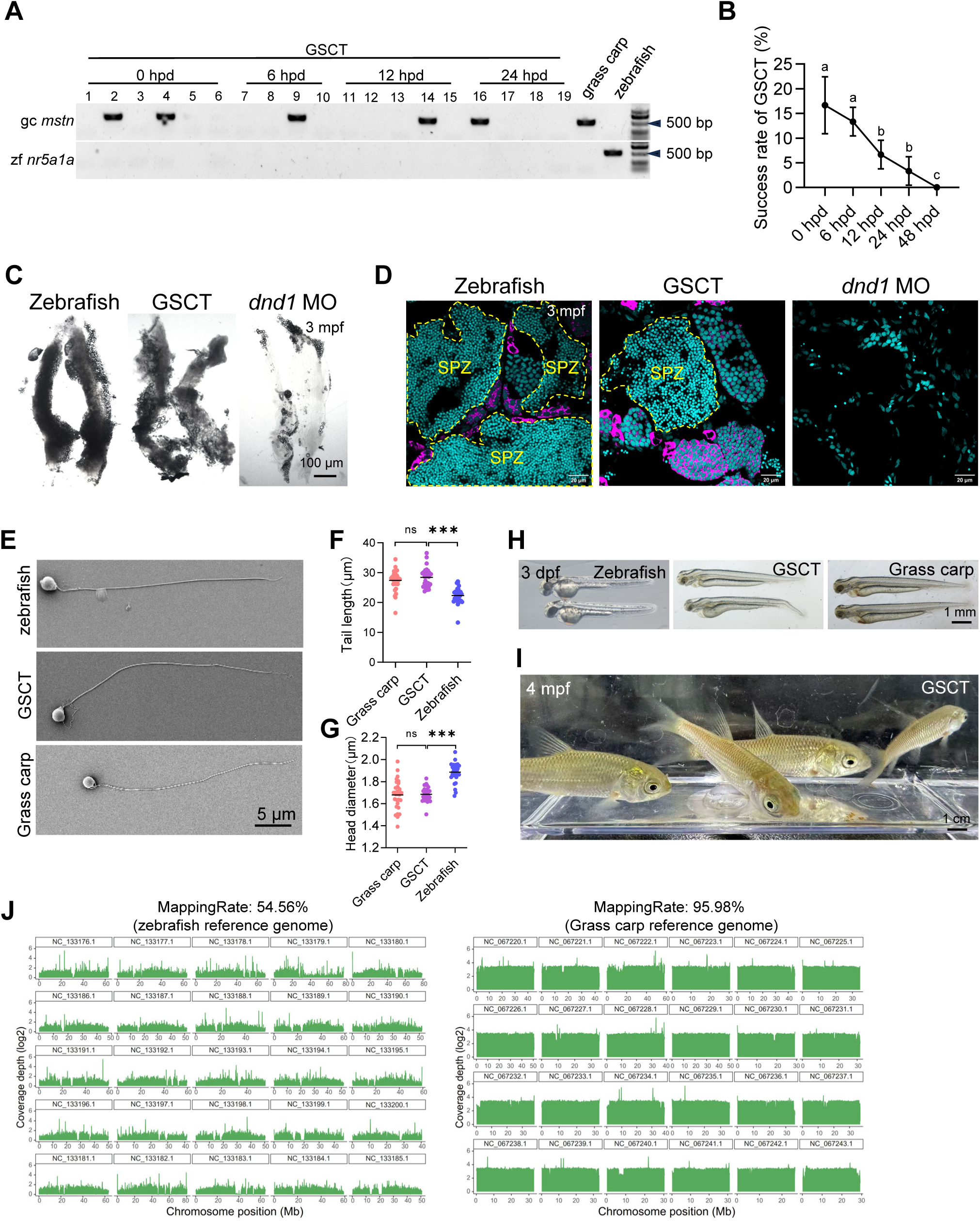
Sperm production and genetic integrity of GSCT zebrafish–derived progeny. A: PCR detection of putative semen samples using grass carp–specific (*gc_mstn*) and zebrafish-specific (*zf_nr5a1a*) primers. B: Proportion of sperm PCR-positive fish (mean ± SEM, N = 3). Different letters indicate p < 0.05 (Tukey–Kramer test). C: Gonadal morphology of wild-type zebrafish, GSCT zebrafish, and dnd1 morpholino-injected zebrafish. D: Immunofluorescence staining of gonads from the indicated groups showing Ddx4-positive germ cells. E: Scanning electron microscopy images of sperm from zebrafish, GSCT zebrafish, and grass carp. F-G: Quantification of sperm head length and flagellum length. ns, not significant; p < 0.05 (Student’s t-test). H: Comparison of larvae at 3 dpf derived from fertilization using GSCT-derived sperm. I: GSCT-derived offspring at 4 mpf. J: Whole-genome resequencing analysis of GSCT-derived larvae.

To further characterize the donor-derived sperm, field-emission scanning electron microscopy (FE-SEM) was employed to compare the morphology of zebrafish sperm, grass carp sperm produced in zebrafish hosts (GSCT sperm), and wild-type grass carp sperm. GSCT sperm exhibited tail lengths and head sizes comparable to those of wild-type grass carp sperm. Compared with zebrafish sperm, GSCT sperm possessed longer flagella and smaller head diameters (Fig. 5E–G), indicating a high degree of morphological similarity to native grass carp sperm. In addition, in vitro fertilization assays were performed to assess the functional competence of GSCT sperm. GSCT sperm successfully fertilized wild-type grass carp eggs, and at 3 dpf, the resulting larvae displayed morphological characteristics indistinguishable from grass carp and clearly distinct from zebrafish larvae (Fig. 5H, I and Supplementary Video 1). Most GSCT-derived offspring developed normally, confirming the full functionality of GSCT sperm.

To further demonstrate that zebrafish can be used to rapidly generate offspring from deceased grass carp, whole-genome resequencing was performed on larvae produced by fertilization with GSCT-derived sperm. Genomic DNA was extracted from larval samples of grass carp, zebrafish, and GSCT-derived F1 offspring. Sequencing was conducted on the Illumina NovaSeq platform, yielding an average paired-end sequencing depth of approximately 30.62× per sample. Raw sequencing data were subjected to quality filtering to remove low-quality reads and adapter contamination. After filtering, each sample yielded an average of approximately 7.65 Gb of high-quality data, with a Q30 value of 98.04%, indicating high sequencing quality. High-quality reads were aligned to the grass carp and zebrafish reference genome, and genome coverage uniformity was assessed based on read depth across genomic regions. When the high-quality reads from GSCT were aligned to the grass carp reference genome, the coverage rate was 95.98%, whereas alignment to the zebrafish reference genome yielded a coverage rate of only 54.56% (Fig. 5J). This indicates that the genome of the larvae fertilized by GSCT-derived sperm is nearly identical to that of grass carp. These results demonstrate that, through germline stem cell transplantation, zebrafish can indeed be used to generate offspring from deceased grass carp.

In summary, by using zebrafish as hosts, we achieved the genetic recovery of deceased grass carp individuals within only three months, providing a rapid and effective strategy for the conservation and recovery of valuable aquatic genetic resources.

## 4. Discussion

Genetic resurrection of aquaculture and endangered fish species represents a crucial strategy for aquatic genetic resource conservation. Here, we establish an ultra-fast genetic recovery platform based on germline stem cell transplantation. Using grass carp as a representative warm-water species, we demonstrate that postmortem germline stem cell viability and transplantation efficiency are tightly linked to tissue freshness and are strongly preserved by low-temperature storage. Transplantation of germline stem cells from deceased grass carp into zebrafish recipients enabled rapid cross-species colonization and differentiation, producing functional donor-derived gametes within three months and generating normally developing offspring. This approach effectively circumvents limitations imposed by body size, long reproductive cycles, and husbandry constraints, offering a rapid and broadly applicable strategy for postmortem genetic rescue.

In this study, we observed that germ cells within the gonads retain biological activity for a substantially longer period after death than surrounding somatic cells (Fig. 1F and 1G), thereby providing a critical time window for genetic rescue of deceased individuals. From a cell biological perspective, gonadal somatic cells typically perform metabolically demanding structural and secretory functions and are therefore highly dependent on continuous energy supply and homeostatic regulation (Schulz *et al*, 2010). Following death, abrupt oxygen deprivation and rapid ATP depletion preferentially trigger apoptotic pathways in these cells, leading to their rapid loss of viability. In contrast, germ cells—particularly germline stem cells—exhibit relatively low metabolic rates, slow proliferation, or even a quasi-quiescent state, and are enriched for stress-resistance and DNA repair mechanisms (Jackson & Finley, 2024; Yan *et al*, 2023; Zhang *et al*, 2022b). These features confer enhanced tolerance to extreme postmortem conditions. Consequently, during the early postmortem period, even when overall tissue freshness declines markedly, a subset of germ cells may retain transplantable and differentiative potential. This intrinsic resilience constitutes the biological foundation for achieving genetic resurrection of deceased fish via germline stem cell transplantation.

Although this study successfully generated functional sperm derived from deceased grass carp individuals, this outcome represents genetic recovery rather than complete individual resurrection in a strict sense. At present, germline stem cell transplantation predominantly yields male gametes, a limitation closely linked to the gonadal microenvironment of the recipient fish (Kikuchi *et al*, 2020; Ren *et al*, 2024). In small model species such as zebrafish, depletion of endogenous germ cells typically biases the gonadal niche toward male differentiation, thereby favoring colonization and spermatogenesis of transplanted germline stem cells while disfavoring oogenesis (Deng *et al*, 2025; Zhang *et al*., 2022a). As a result, even when donor cells originate from females or undifferentiated individuals, the type of gamete ultimately produced remains strongly constrained by the recipient microenvironment. Achieving true genetic resurrection of deceased individuals will therefore require the identification or engineering of recipient systems (Okutsu *et al*, 2006) capable of supporting oocyte development, as well as targeted modulation of gonadal niche factors and sex determination pathways to overcome this inherent male bias.

Germline stem cell transplantation is not the sole route to genetic resurrection. For individuals from which transplantable germ cells can no longer be recovered after death, alternative strategies such as somatic cell nuclear transfer (Bail *et al*, 2010; Hossein *et al*, 2021; Luo *et al*, 2011) combined with induced primordial germ cell (iPGC) technologies provide important complementary options (Mullins, 2024; Wang *et al*., 2023). Although somatic nuclear transfer in fish continues to face challenges including low developmental efficiency and incomplete epigenetic reprogramming (Depincé *et al*, 2021; Rouillon *et al*, 2019), it remains valuable in specific species and rescue scenarios. Meanwhile, reprogramming somatic cells into iPGCs followed by transplantation into suitable hosts offers a theoretically viable path for genetic transmission when endogenous germ cells are entirely lost. Looking forward, the strategic integration of germline stem cell transplantation, nuclear transfer, and iPGC-based approaches—selected according to postmortem freshness, available cell types, and species-specific characteristics—may enable the construction of a tiered and flexible genetic resurrection framework. Such a system would maximize the success rate and applicability of genetic resource rescue in aquaculture and endangered fish species conservation.

## Acknowledgments

We sincerely thank Luyuan Pan and Linglu Li from the China Zebrafish Resource Center (CZRC) for their technical assistance *in vitro* fertilization, Fang Zhou and Guangxin Wang from the analytical and testing center of Institute of Hydrobiology, CAS for their support with confocal imaging, and Yuan Xiao and Zhengfei Xing from the analytical and testing center of Institute of Hydrobiology, CAS for their assistance with sperm imaging. This work was supported by the STI2030—Major Projects (2023ZD04055), the Strategic Priority Research Program of Chinese Academy of Sciences (CAS) (XDB0730300), the National Natural Science Foundation of China (32025037 and 32403020), Natural Science Foundation of Hubei Province (2025AFA053), Natural Science Foundation of Wuhan (2024040701010069), Science and Technology Special Fund of Hainan Province (ZDYF2024XDNY256), and State Key Laboratory of Breeding Biotechnology and Sustainable Aquaculture (2024BBSA01).

